# Reward & Imitation in Social Conformity

**DOI:** 10.64898/2026.01.15.699770

**Authors:** Garrett Mauter, Mimi Liljeholm

## Abstract

Normative social conformity has been proposed to elicit a hedonic reward signal that is dissociable from informational inferences about decision outcomes. If present, such a signal should reinforce not just the decision that preceded it, but also any incidentally co-occurring stimulus features. Alternatively, normative conformity might reflect a non-hedonic imitation algorithm. Across two studies (n=359) we used a non-deceptive multi-participant gambling task in which trial-by-trial information was provided about the selections and monetary payoffs of two other participants facing the same, recurring, options in real time. Consistent with both accounts, and contrary to mere monetary maximization, the probability of staying with a losing option increased with the degree of decision unanimity. However, contrary to the social reward hypothesis, only monetary payoffs modulated the valence of incidental gambling stimuli. A prosocial framing did not significantly alter this pattern of results, which favors an imitative over a hedonic account of normative social conformity.

## Introduction

Social conformity – abandoning one’s judgement in favor of aligning with a group majority – has been argued to reflect two distinct processes: the *informational* use of other’s responses as evidence for decision outcomes, and an intrinsic drive towards aligning with apparently *normative* behavior^1^. Informational motivations are likely to dominate when objective feedback about outcomes is lacking^2–6;^ conversely, majority alignment of subjective evaluative judgements, e.g., about the attractiveness of a face or appetitiveness of a food, is commonly attributed to normative conformity^7–10^. Intriguingly, alignment of evaluative judgements has been shown to correlate with BOLD activity in brain regions heavily implicated in reward processing^7, 9, 11^, and such demonstrations have been taken to suggest that normative conformity elicits a generic reward signal^12–15^, commonly formalized as a Temporal Difference (TD) error^12–18^. Here, we pit this ‘reward surrogate’ explanation against an alternative account, positing that subjective evaluations towards an ostensible norm might reflect a valence-neutral action-copying algorithm, previously considered as a basis for imitative observational learning^16, 19–23^. We devised a multi-participant gambling task to test dissociable predictions of these two accounts.

The task was administered online, with individuals participating in groups of three. On each trial, all participants in a group were presented with a pair of abstract shapes (drawn from a set of six), prompted to select one of the two, and then shown the monetary payoffs of both options on that trial, as well as the selections made by the other two participants (see Figure 1A). To further emphasize the monetary difference between trial outcomes, the lower of the two payoffs was reduced to zero following the initial monetary feedback. Thus, six outcome scenarios of interest were generated by combining levels of social alignment (i.e., with one, both, or neither of the other participants). with a monetary gain vs. opportunity cost. To ensure variability in decision unanimity, both options presented on a given feedback trial were drawn randomly from the *same* reward distribution and modified to ensure at least a $0.1 difference. While this generated necessary uncertainty about the ‘correct’ choice on a given trial, the trial-based feedback categorically ruled out the validity of other participant’s decisions as reliable sources of information about decision outcomes.

**Figure 1:**
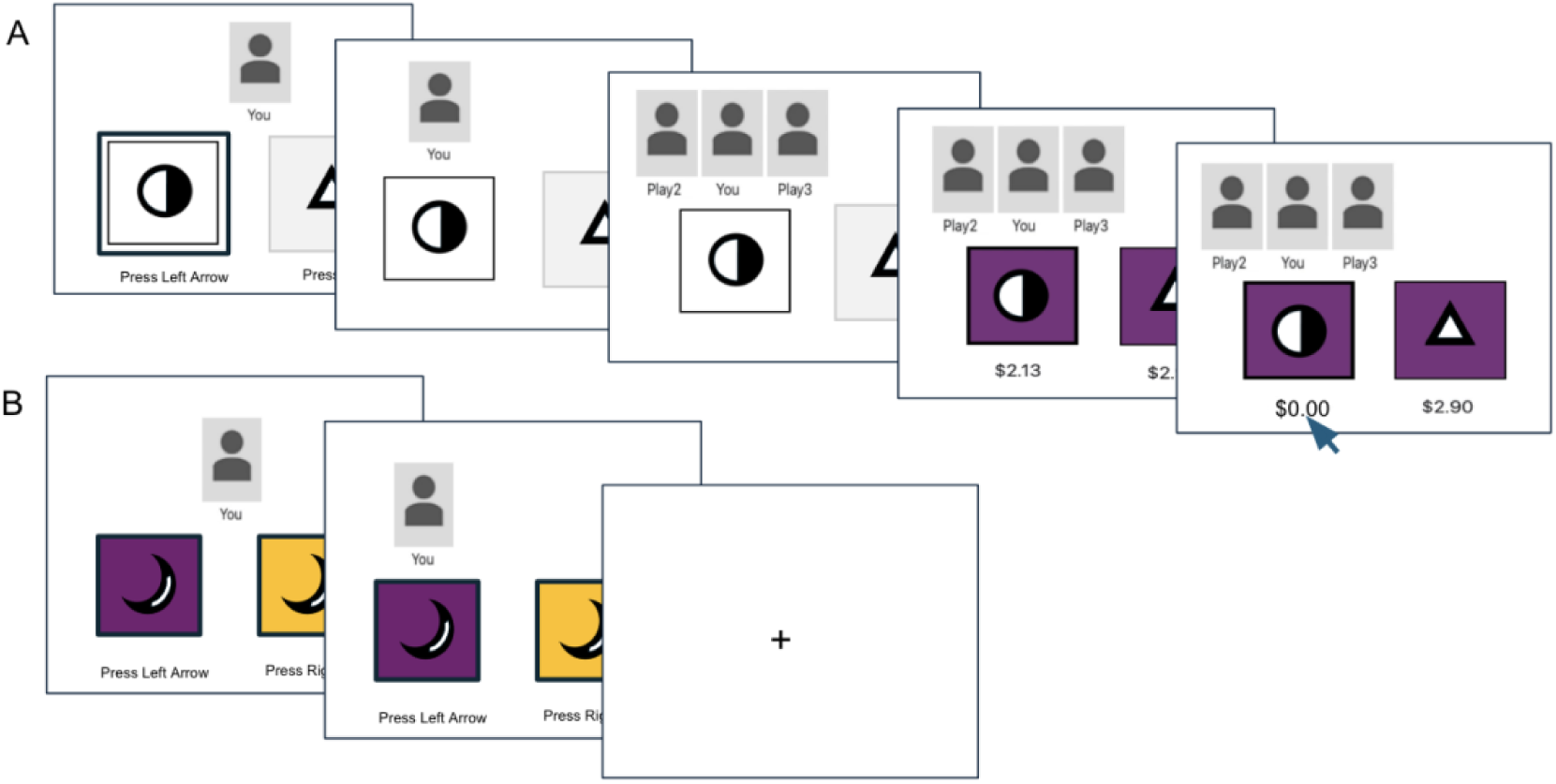
Trial Illustration. **A**. On gambling trials, participants are presented with two gambling options (abstract black and white shapes). Upon selection, the participant’s avatar, initially positioned top center, moves to align with the chosen option. On the subsequent feedback screen, the decisions of the other two participants are likewise indicated by the alignment of their avatars, followed by the display of monetary payoffs beneath each option and a color change applied to *both* options, with each color corresponding to one of six outcome scenarios (see text). Finally, the lower of the two payoffs is set to zero (arrow), to emphasize the monetary difference between trial outcomes. **B**. On transfer trials, different-color options with novel shapes are presented but no social or monetary feedback is provided following selection.

In addition to social and monetary feedback, a condition-specific background color was applied to *both* options on feedback screens, with a unique and counterbalanced color, orthogonal to gambling options and decisions, repeatedly paired with each of the six feedback scenarios. To assess a generic reward signal elicited by monetary gain and/or social alignment, intermittently occurring transfer trials solicited a choice between color options that had been paired with consensus vs. dissent, respectively, on preceding trials (see Figure 1B). Critically, to preclude learning, no feedback about monetary or social trial outcomes were provided on transfer trials, some of which also solicited selection between gambling (i.e., shape) options associated with different reward distributions.

We specified three learning algorithms that differed with respect to their treatment of other’s decisions and assessed the relative fit of these models to choice behavior and evaluative judgements: A Baseline model that only considers monetary outcomes as rewarding; a Social learner that treats consensus as a surrogate reward; and an Imitation model, that treats only monetary outcomes as rewarding, but that strived to copy observed decisions in conjunction with reward maximization. While all three models predict value estimation based on monetary payoffs, only the Social and Imitation accounts predict a tendency to repeat others’ decisions, and only the Social learner predicts a transfer of valence to incidental stimuli that covary with majority alignment.

## Study 1

### Methods

#### Participants

One hundred and twenty participants (58 females, mean age = 33.2 ± 10.9) were recruited and completed the study on Prolific (www.prolific.com) for monetary payment. One participant was excluded for pressing the left key on 97% of the trials. All participants gave informed consent and the Institutional Review Board at the University of California, Irvine approved the study. All aspects of the study conformed to the guidelines of the 2013 WMA Declaration of Helsinki. The sample size was set to match a methodologically similar multiplayer task manipulating social information and objective performance^24^.

#### Task & Procedure

Upon joining the study and providing consent, participants were asked to wait until the group totaled three individuals (average wait time was ∼3 minutes). To emphasize the social element, participants were asked at the start of the study to enter individual ‘player names’ to be displayed beneath each player’s icon, so that they could be identified by the other participants throughout the task. They were instructed that they would be selecting between various gambles with different monetary payoffs, individually but in parallel, and that, on each gambling trial, they would receive feedback about each other’s decisions and monetary outcomes as well as their own. All three participants had to submit their selection for a trial to proceed, and all decisions had to be submitted within 10 seconds, or the trial would be canceled. Before the gambling phase, participants were asked to rate the pleasantness of each of 6 distinct colors, to be used as contextual stimuli, on a scale from 0 (not at all pleasant) to 10 (extremely pleasant).

Critically, the two (of six) distinct shapes appearing as options on each gambling trial had been randomly assigned at the start of the experiment to *the same* of two reward distributions with ‘high’ (µ=$4.5±$1.5) and ‘low’ (µ=$2.0±$1.5) means respectively. Once all participants had made their selection, monetary payoffs for both options were drawn from the shared distribution, one of which was modified if needed to ensure at least a $0.1 difference. The payoffs were then displayed on the screen, together with the group alignment, and the lower of the two was set to $0, in order to emphasize the Win vs. Loss outcome of the trial. One shape from each distribution was included to provide variability, in order to prevent fatigue and disengagement, and thus occurred with about half the frequency of the other four shapes. At the end of the study, 3 trials were randomly drawn from all feedback trials, and each participant received the sum of their individual earnings on those 3 trials.

The group alignment on each gambling trial (see Figure 1A) occurred naturally as a consequence of participants’ choices, without experimental deception or manipulation, and independently of monetary outcomes, since both options were associated with the same monetary reward distribution. In contrast, a unique, and participant-specific, outcome color was applied to *both options* following choice, that depended on the particular combination of monetary and social outcomes, specifically, the following conditions: 1. Win (selected the greater payoff option) and Full (both of the other two participants selected the same option), 2. Win and Partial (one of the other two participants selected the same option), 3. Win and Dissent (neither of the other two participants selected the same option), 4 Lose (selected the lesser payoff option, resulting in a $0 trial outcome) and Full, 5. Lose and Partial, and 6. Lose and Dissent. To reduce the color counterbalancing load, two colors were randomly assigned to the Partial conditions (for which predictions are somewhat equivocal and which necessarily occur about twice as often as Full Consensus and Dissent), while a separate set of four primary colors were counterbalanced across the remaining four outcome states.

In addition to gambling trials, intermittent ‘transfer’ test trials solicited a selection between options displaying the same, otherwise not shown, shape but with *different* background colors, that were respectively paired with consensus vs. dissent, and with wins vs. losses, on gambling trials (see Figure 1B). Critically, no social or monetary feedback was provided on these trials, to prevent learning effects. To confirm that participants were tracking monetary payoffs, additional transfer trials forced selection between shape options associated with different reward distributions, again without providing any feedback.

#### Computational Modeling & Statistical Analyses

We specified three error-driven learning rules that differed with respect to their treatment of other’s decisions and assessed the relative fit of those models to choice behavior. First, a Baseline model considered only monetary outcomes as rewarding;

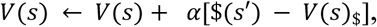

where *V(s)* is the value estimate for stimulus, *s* (i.e., a gambling option or background color), 𝛼 is a learning rate parameter and $*(s’)* is the monetary payoff paired with option *s*. Second, a Social Reward learner that treats majority alignment as a surrogate reward that scales with majority size:

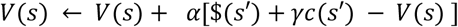

where *c(s’)* is the conformity outcome, reflecting alignment with 2, 1 or 0 other players, and γ is a free parameter estimating the subjective utility of conformity. Finally, we specify an *Imitation* model that treats only monetary outcomes as rewarding, but that strived to copy observed decisions in conjunction with reward maximization:

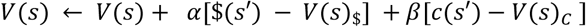

where 𝛽 is a learning rate parameter for action copying.

Note that, in addition to variations in the frequency of gambling options, the task necessarily generates more partial consensus trials than full consensus or dissent trials. We added a UCB1 term to value estimates to account for different event frequencies in both gambling options and incidental features and the resulting value was passed to a softmax rule with a free noise parameter that generated decision probabilities. Models were fit to behavioral data by minimizing the negative log likelihoods and the Akaike Information Criterion (AIC) was used for model comparisons. Parameter recovery was assessed by simulating 1000 players, each receiving payoffs and social information drawn from randomly selected participant data, with randomly generated parameter values, from the range used in model fitting, and with Pearson’s correlation coefficients estimating recovery.

We use planned comparisons and repeated measures Analyses of Variance (ANOVAs) to assess the influence of decision unanimity on behavioral gambling and transfer test trials. We use the term ‘*es*’ throughout to indicate effect sizes for t- (Cohen’s d) and F- (partial eta squared) tests. All tests are two-tailed.

#### Trait Assessments

In addition to the gambling task and pleasantness ratings, participants completed surveys assessing individual differences in Individualism and Collectivism ^25^, Narcissism ^26^, and Social Anxiety ^27^. To ameliorate completion load, these were administered between participants, with each participant receiving either the Individualism and Collectivism surveys or the Narcissism and Social Anxiety surveys, in counterbalanced order. These questionnaires were administered at the end of the experiment for exploratory purposes and are noted here only for completeness.

## Results

With respect to model performance, AIC scores, penalizing for free parameters, were lower, indicating better performance, for the Imitation model than for both the Social, *t*(118)=-2.708, *p*<0.01, *se*=-0.25, 95%, CI [-6.94, -1.08], and Baseline, *t*(118)=-3.43, *p*<0.001, *se*=-0.31, 95% CI [-8.03, -2.15], models. Parameter recovery was excellent across models, for gambling (*r>*0.99 & *p<*0.001) and contextual (*r>0.91* & *p<*0.001) learning rates, decision noise (*r>*0.97 & *p<*0.001), and the subjective utility of social reward/imitation learning rate (*r>*0.99 & *p<*0.001).

We operationalized social conformity as the tendency to stay with a gambling option given the unanimity with which it was selected on the previous trial, if repeated (regardless of left/right positions on the screen). Mean behavioral stay probabilities as a function of decision unanimity and monetary payoffs, are plotted in Figure 2A, together with corresponding choice proportions from model simulations (Figs. 2C-E). Planned comparisons confirmed that, as predicted by both the Social and Imitation models, but not the baseline (monetary) model, the probability of staying with an option that had incurred a loss (i.e., $0 payoff) on the previous trial was significantly greater for unanimous, Consensus, decisions than for Dissent decisions, that deviated from both other participants; *t*(110) = 2.30, *p=*0.02, *es*=0.22, 95% CI=[0.01, 0.15]. A repeated measures Analysis of Variance (ANOVA) with Consensus & Payoff as within-subject factors, revealed significant effects of Consensus [F(2, 214)=3.50, *p*=0.03, *es=*0.03] and Payoff [F(1, 107)=9.73, *p*=0.002, *es=*0.08], but no interaction (p=0.84).

**Figure 2:**
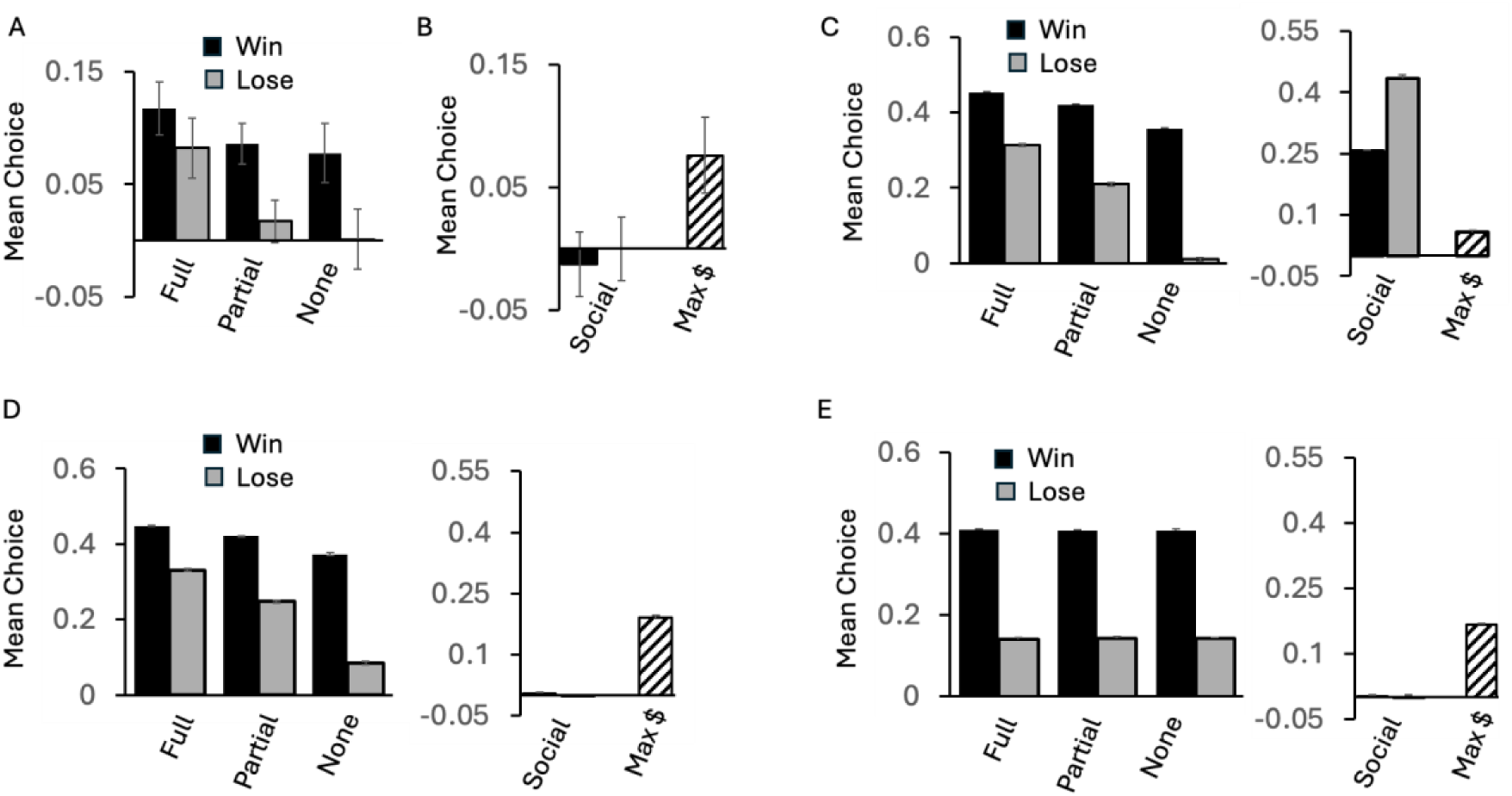
Behavioral & Simulated choice proportions with chance performance subtracted, from Study 1. A) The probability of repeating selection of an option that occurred on the previous trial, as a function of decision unanimity (full, partial, or none) & monetary (win vs. loss) outcome on that previous trial. B) Choice proportions on transfer trials without feedback. Left two bars show preference for contextual stimuli (background colors) paired with Consensus over Dissent (i.e., full unanimity vs none), in Win and Lose conditions respectively. Rightmost bar shows degree of preference for gambling options associated with High vs. Low mean reward distributions. C), D), and E) respectively show corresponding predictions by simulated Social Reward, Imitation, and Baseline learners. Error bars = SEM.

While both the Imitation and Social Reward learners predict the influence of consensus on stay probabilities, the Social Reward model alone predicts that majority alignment will elicit a generic reinforcement signal that transfers not only to relevant decision variables, but also to any incidentally occurring contextual features. We assessed such reinforcement using transfer trials without feedback, in which the two gambling options had the same shape, but *different* colors, one being deterministically associated with unanimous, Consensus, decisions and the other with Dissent from both other participants. Mean choice preference for Consensus over Dissent, for monetary Win and Loss colors, are illustrated on the left side of Figure 2B. Planned comparisons confirmed that there was no significant Consensus preference for either Win (p=0.68) or Loss (p=0.50) colors. In contrast, as shown on the right side of Figure 2B, transfer trials that assessed a preference for gambling options associated with greater monetary payoffs revealed a clear effect; t(118)=3.07, *p=*0.003, *es=*0.28, 95% CI=[0.03, 0.12].

Finally, a consensus-induced change in the valence of condition-specific background colors was assessed using pleasantness ratings solicited before and after the gambling session. The difference between pre-and post-gambling ratings, as a function of social and monetary contingencies, is plotted in Figure 3A. A repeated measures ANOVA performed on the difference between pre-and post-gambling ratings, with Consensus and Win/Lose Payoff as factors, revealed a significant main effect of Payoff, F(1, 118)=5.34, *p*=0.023, *es=*0.04, indicating that participants affect did indeed change for contextual features that coincided with monetary gain, but no significant effect of Consensus and no interaction (*p>*0.29), again contrary to the social reward hypothesis.

**Figure 3.**
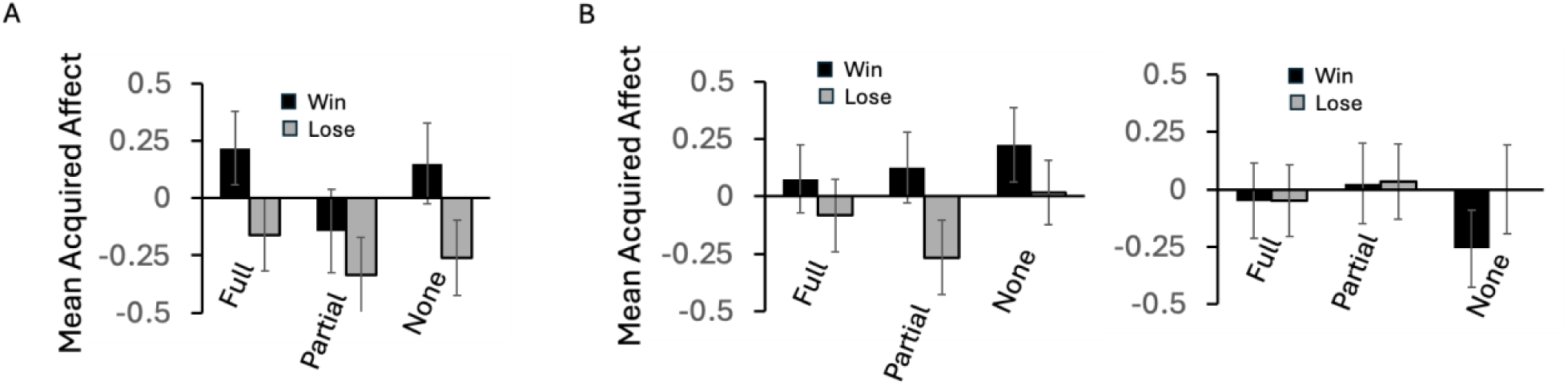
Changes in evaluative judgements. The difference between pleasantness ratings of contextual color stimuli obtained before and after the gambling phase, plotted as a function of their pairing with a particular level of group alignment (full, partial, or none) and with Win vs. Lose monetary outcomes where a loss indicates a $0 trial payoff. A. Results from Study 1. B. Results from the Control (left) and Charity (right) groups in Study 2. Error bars = SEM.

### Interim Discussion

In Study 1 we found clear evidence of normative conformity, in that participants were significantly more likely to stay with a losing option that repeated on the subsequent trial if it had been unanimously selected than if both other participants had selected the other alternative. We also found clear evidence of monetary reward maximization, and of changes in affect associated with contextual stimuli based on monetary payoffs; however, critically, there was no evidence of changes in contextual affect based on Consensus vs. Dissent. One limitation of these results is that monetary reinforcement of contextual features was assessed using evaluative judgements only, while social reinforcement, pivotal for ruling out a social reward surrogate, was assessed using both evaluative judgements and transfer trials during gambling. We address this limitation in Study 2.

Another potential weakness of Study 1 is that it may be lacking conditions that are necessary for conformity to become rewarding. Although previous claims about the intrinsic reward of social conformity have not posited any contextual constraints^6, 7, 13, 17, 28^, a large number of studies have focused on conformity in prosocial contexts^29–32^. In Study 2, one group of participants made all gambling decisions knowing that any monetary earnings would be donated to a charity of their choice, rather than benefit themselves financially. We hypothesized that the well-documented peer pressure mediating prosocial decisions might make majority alignment more rewarding, shifting the evidence towards the ‘conformity as reward’ account.

## Study 2

### Methods

#### Participants

Two-hundred and forty participants (114 females, mean age = 41.6 ± 11.2) were recruited and completed the study on Prolific (www.prolific.com) for monetary payment. The sample size was set to match that of Study 1, with 120 subjects randomly assigned to each of two groups, a Control group and a Charity group, and with no exclusions. All participants received a base payment of $10 for participation, with additional performance-based bonuses benefiting either the participant (Control group) or a charity of their choice (Charity group). All participants gave informed consent and the Institutional Review Board of the University of California, Irvine, approved the study. All aspects of the study conformed to the guidelines of the 2013 WMA Declaration of Helsinki.

#### Task & Procedure

The task and procedures were identical to Study 1, except that in one group participants received instructions that all earnings would be donated to a charity; once the gambling phase was completed, participants in this group were prompted to select a charity from a 40-item list of American and international charities, to which their bonuses were subsequently donated. Moreover, contrary to Study 1, no trait-assessment questionnaires were administered and background colors were fully counterbalanced across social conditions, including partial consensus.

## Results

As in Study 1, AIC scores were lower, indicating better performance, for the Imitation model than for both the Social, *t*(119)=-2.30, *p*=0.02, *es=*0.21, 95% CI [-10.91, -0.85], and Baseline, *t*(118)=-5.43, *p*<0.001, *es=*0.50, 95% CI [-17.16, -7.99], models. Mean behavioral stay probabilities as a function of decision unanimity and monetary payoffs are plotted in Figures 4, for the Control (A) and Charity (C) group respectively. Planned comparisons, collapsing across groups, revealed that the probability of staying with an option that had incurred a loss (i.e., $0 payoff) on the previous trial was significantly greater for unanimous decisions than for decisions that dissented from both other participants; *t*(230) = 2.23, *p=*0.027, *es*=0.15, 95% CI=[0.01, 0.12]. A mixed ANOVA performed on behavioral stay probabilities, with Group as between-subjects factor and Consensus and Monetary Payoff as within-subject factors, revealed significant effects of Consensus [F(2, 444)=6.13, *p*=0.002, *es=*0.03] and Payoff [F(2, 444)=21.19, *p*<0.001, *es=*0.09], but no other main effects or interactions (p>0.39).

**Figure 4:**
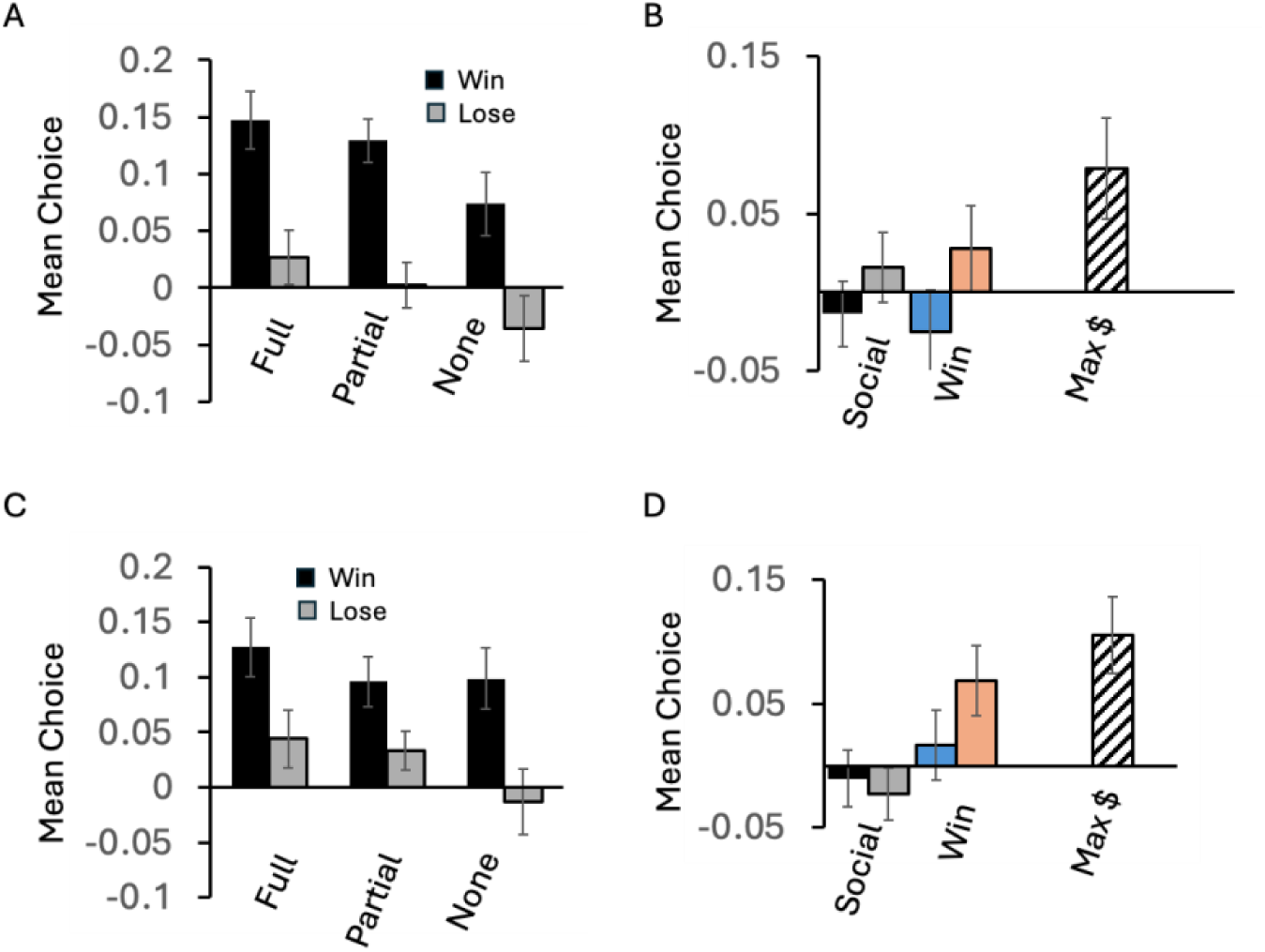
Behavioral results from Study 2. A) Stay Probabilities (deviation from chance) as a function of Social & Monetary Outcomes in the Control group. B) Choice proportions on transfer trials in the Control group: Leftmost two bars show degree of preference for cues paired with Full Consensus (over Dissent), for monetary Win and Lose cues respectively. Blue and red bars show the degree of preference for cues paired with monetary Win (over loss) outcomes, for Consensus and Dissent cues respectively. Rightmost single bar shows degree of preference for gambling options associated with High vs. Low mean reward distributions. C & D) Corresponding results in the Charity group. Error bars = SEM.

As in Study 1, despite apparent conformity in gambling decisions, no significant preference was found for background colors paired with full Consensus over those paired with Dissent (leftmost two bars in Figures 4B and 4D), in either Win (0.78) or Loss (*p=*0.57), conditions, again suggesting the absence of a generic reward signal elicited by group alignment. In contrast, as shown with blue and red bars in Figures 4B and 4D, there was a clear preference for background colors associated with monetary Win (over Loss) outcomes, though interestingly this preference did not emerge for Win colors associated with Consensus (*p=*0.85), only for Win colors associated with Dissent; t(239)=2.64, *p=*0.009, *es=*0.17, 95% CI=[0.01, 0.09]. As in Study 1, the strongest transfer effects emerged for choices between gambling options drawn from different monetary reward distributions (never pitted against each other on feedback trials). A mixed ANOVA revealed a significant effect of reward distribution [F(2, 238)=26.37, *p*<0.001, *d=*0.10], but no significant effect of group or interaction (p>0.5)

Unlike choice probabilities on feedback and transfer trials, evaluative judgements of contextual cues looked different in the Charity group (right side of Figure 3B) from both the Control group (left side of Figure 3B) and the Study 1 group (Figure 3A). Specifically, whereas in the Control group, acquired affect again appeared to be solely based on monetary outcomes, those in the Charity did not vary consistently with either monetary or social outcomes, resulting in a just-significant Payoff-by-Group interaction; F(1, 238)=3.89, *p*=0.050, *es=*0.02. No other main effects or interactions were significant (*p>*0.26) .

## General Discussion

Across two studies, we assessed whether normative social conformity is accompanied by a hedonic reward signal^7–10^ that, if present, should reinforce antecedent decisions as well as contextual stimulus features^33, 34^. We contrast this hedonic reward account with a non-hedonic action-copy algorithm, previously argued to support imitative observational learning^22^. Our non-deceptive, real-time, multi-participant economic choice task pitted monetary gain against group alignment, in order to test divergent predictions by these approaches. Specifically, we assessed whether valence elicited by social alignment transferred to incidental stimuli (according to basic principles of reinforcement learning) and whether such alignment-based rewards were integrated with monetary payoffs to compute expected utility on a shared scale. Contrary to the frequent interpretation of behavioral and neural correlates of conformity as reflecting hedonic reward^7–10^, we found that an Imitative error-driven learner accounted better for choice behavior, with respect to the preference for explicitly defined gambling options as well as incidental (i.e., contextual) stimuli.

Three aspects of our task constitute significant departures from previous work in this vein: First, we are allowing group alignment to evolve naturally and dynamically in a non-deceptive decision environment, where real-time behavior is collected from multiple participants. This is in contrast to commonly used fictional ‘norm’ decisions (e.g., mean judgements by ostensible previous participants^7–10, 17, 35^). Second, we provide immediate feedback about the trial-level accuracy of others’ decisions, a feature that is virtually absent in conformity research, despite the obvious use of decision-contingent outcomes in real-world judgements. When provided at the trial level, information about the outcomes of observed decisions should shape rational informational conformity, but not normative conformity behavior which, by definition, is void of informational content^1^. Finally, unlike previous work using reward learning to explain normative social conformity, we separate the postulated social reinforcement signal from that of making a reward-maximizing (or accurate, or norm-aligned) decision, by assessing the transfer of valence to contextual stimuli.

We found a clear influence of both decision unanimity and monetary outcomes on stay probabilities in gambling decisions, as well as independent evidence of monetary, but not social, reinforcement of gambling options and contextual features on transfer trials. These results favor an imitative account over the idea of group alignment as a reward surrogate. Notably, arguments favoring social reward often rest on neuroimaging data demonstrating that an individual’s adjustment of subjective evaluations towards a group norm is predicted by BOLD activity in the ventral striatum and ventromedial prefrontal cortex ^7–10^ – areas heavily implicated in the detection and integration of reward information ^12, 36–40^. However, these brain regions are responsive to a range of sensory properties and conditions: For example, the ventral striatum also responds to a stimulus’ aversiveness ^41–43^ and novelty ^44–47^ and the ventromedial prefrontal cortex has been shown to encode non-valanced social information, such as self-relevance^48^. Thus, behavioral adjustments of subjective evaluative judgements towards a highlighted norm might reflect a more general, potentially non-hedonic, form of error correction, a possibility that is supported by the current results.

In Study 2, we assessed whether pro-social decision making, often used to assess conformity^29–32, 49^, would enhance the perceived utility of group alignment. As in Study 1, we found no evidence of a preference for contextual stimuli associated with consensus decisions, in either the prosocial or self-benefiting group. However, we did find a significant preference for contextual stimuli associated with monetary wins that was specific to dissent outcomes, indicating a potential integration of social and monetary variables. This asymmetry might reflect a tacit assumption on the part of participants that consensus decisions entailed splitting the payoff across group members, and thus that such trials were *less* valuable in terms of monetary profit. While the existence of such a bias could overshadow an intrinsic utility of consensus on win trials, a clear preference should still have been apparent on loss trials. Moreover, even a shared profit on win trials should result in a preference relative to the loss ($0) outcome, particularly for payoffs drawn from the relatively high reward distribution. We conjecture instead that unanimous decisions either distracted from the processing of monetary information or detracted from the intrinsic utility of ‘being right’, a significant motivational factor in its own right, not explicitly addressed here. Future work is needed to arbitrate between these possibilities.

In conclusion, our findings highlight commonalities between model-free imitative learning and normative social conformity. This mapping can be extended to associated constructs of informational social conformity (treating others’ decisions as indicative of decision outcomes^1^) and emulative observational learning (reproducing a goal state rather than copying an action^39^), perhaps with a common formalization as inverse reinforcement learning, previously applied to emulation^23^. An important implication of such mappings is that they generate divergent predictions about an acquired behavior based on which learning strategy was employed during acquisition - in particular, model-free acquisition leads to less flexible behavior^50^. We will probe these questions in future work, in order to better characterize the motivational and cognitive bases of social transmission.

